# BamA and BamD are essential for the secretion of trimeric autotransporter adhesins

**DOI:** 10.1101/2020.08.14.252015

**Authors:** Jessica L. Rooke, Chris Icke, Timothy J. Wells, Amanda E. Rossiter, Douglas F. Browning, Faye C. Morris, Jack C. Leo, Monika S. Schütz, Ingo B Autenrieth, Adam F. Cunningham, Dirke Linke, Ian R. Henderson

## Abstract

The BAM complex in *Escherichia coli* is composed of five proteins, BamA-E. BamA and BamD are essential for cell viability and are required for the assembly of β-barrel outer membrane proteins. Consequently, BamA and BamD are indispensable for secretion via the classical autotransporter pathway (Type 5a secretion). In contrast, BamB, BamC and BamE are not required for the biogenesis of classical autotransporters. Recently, we demonstrated that TamA, a homologue of BamA, and its partner protein TamB, were required for efficient secretion of proteins via the classical autotransporter pathway. The trimeric autotransporters are a subset of the Type 5-secreted proteins. Unlike the classical autotransporters, they are composed of three identical polypeptide chains which must be assembled together to allow secretion of their cognate passenger domains. In contrast to the classical autotransporters, the role of the Bam and Tam complex components in the biogenesis of the trimeric autotransporters has not been investigated fully. Here, using the *Salmonella enterica* trimeric autotransporter SadA and the structurally similar YadA protein of *Yersinia* spp., we identify the importance of BamA and BamD in the biogenesis of the trimeric autotransporters and reveal that BamB, BamC, BamE, TamA and TamB are not required for secretion of functional passenger domain on the cell surface.

**Importance:** The secretion of trimeric autotransporters (TAA’s) has yet to be fully understood. Here we show that efficient secretion of TAAs requires the BamA and D proteins, but does not require BamB, C or E. In contrast to classical autotransporter secretion, neither trimeric autotransporter tested required TamA or B proteins to be functionally secreted.

## INTRODUCTION

The Type 5 secretion system (T5SS) is the most abundant protein secretion system in Gram-negative bacteria (1-3). Recently, the T5SS has been divided into five sub categories; Type 5a (classical autotransporters), Type 5b (two-partner secretion), Type 5c (trimeric autotransporters), Type 5d (fused two-partner) and Type 5e (inverted autotransporters) (4-6). All sub-classes of the T5SS have the following domains: a signal sequence, secreted passenger domain and a β-barrel translocation domain. However, the exact composition, order and size of these domains differ between sub-classes (5, 7, 8).

Secretion via the T5SS has mostly been studied in the T5aSS. In this system, the autotransporter protein contains a signal sequence, a passenger domain, and a ca. 300 amino acid β-barrel translocation domain. The signal sequence directs translocation across the inner membrane via the SecYEG translocon (9-11). Once in the periplasm, the signal sequence is cleaved (12). Periplasmic chaperones, such as SurA and DegP, assist in the delivery of the remaining protein to the outer membrane β-barrel Assembly Machinery (Bam) complex (13-15). This complex facilitates the insertion of the C-terminal translocation domain into the outer membrane as a monomeric β-barrel (12). The β-barrel domain of the T5aSS initiates translocation of the passenger domain to the cell surface and once the passenger domain is located extracellularly it adopts a folded conformation (16). Whilst the precise mechanism of passenger domain translocation is yet to be elucidated, the Bam complex is integrally involved in the secretion process (17-19).

In *Escherichia coli* the Bam complex is comprised of five proteins, BamA-BamE (18, 20, 21). BamA is an essential outer membrane protein (22, 23). Four periplasmic lipoproteins, BamB-BamE, form a non-covalent interaction with BamA to create the complex (24, 25). Like BamA, BamD is essential for cell viability (26). The remaining components are not critical for viability but loss of these components compromises the integrity of the outer membrane and the biogenesis of some outer membrane proteins (26). Recent studies assessing the contribution of individual BAM complex proteins to the biogenesis of autotransporters have shown that BamA and BamD are essential for the secretion of the classical autotransporters from *E. coli* (20). In contrast, BamB, BamC and BamE were not essential for the secretion or folding of these proteins (20). Additionally, we have previously reported that the Translocation Assembly Machinery (Tam) complex was also required for efficient secretion of a T5aSS protein (27). The Tam complex consists of the integral outer membrane protein TamA and the integral inner membrane protein TamB; TamA shares structural similarity with BamA. Recent studies have demonstrated that the Tam complex facilitates efficient assembly of both Type I fimbriae and Type 5e autotransporters (28, 29). However, the precise contribution of the Tam complex to the biogenesis of all members of the T5SS is yet to be elucidated.

In contrast to the T5aSS, the mechanisms of secretion via the other subclasses are less understood. Proteins belonging to the T5cSS, or the Trimeric Autotransporter Adhesin (TAA) family, possess key differences when compared to members of the T5aSS (30, 31). TAAs are defined by the presence of a short 70-100 amino acid C-terminal translocation domain, and thus the protein must form a homotrimer in order to generate a complete β-barrel (32-35). Consequently, unlike the passenger domains of the T5aSS, which predominantly adopt a monomeric β-helical conformation, the TAA passenger domains form a trimeric structure composed of oligomeric coiled-coil regions interspersed with distinct head and neck motifs (31). The molecular organisation of these proteins presents logistical challenges for the bacterium. For example, the trimer must assemble in a manner that is consistent with the secretion of large passenger domains (up to 5 MDa) to the bacterial cell surface; at the outer membrane each monomer must be maintained in a conformation that is consistent with pairing to its sister monomers; and monomers must be inserted into the outer membrane whilst being prevented from pairing with the nascent β-strands of non-partner proteins. Previously, using the archetypical and widely studied TAA protein, YadA from enteropathogenic Yersiniae, we demonstrated that BamA was essential for the assembly of a folded functional trimer into the outer membrane (36). Although the Tam complex has been implicated during the biogenesis of some T5aSS proteins, loss of TamA did not affect the outer membrane accumulation or folding of two TAA proteins (37). Due to the complexity of T5cSS, we hypothesised that the other components of the Bam complex and/or the components of the Tam complex would play important roles in co-ordinating the assembly of the TAA homotrimer into the outer membrane. Here we tested this hypothesis by determining the impact that loss of each Bam and Tam component had on the assembly and function of two TAAs, YadA from *Yersinia* spp., and SadA from *Salmonella enterica*.

## MATERIALS AND METHODS

### Bacterial strains and growth conditions

The strains and plasmids used in this study are listed in Table 1. All strains were grown in lysogeny broth (LB) with appropriate antibiotics unless otherwise stated. Strains that required antibiotics were supplemented with either 100 µg/ml ampicillin or 50 µg/ml kanamycin. The BamA and BamD depletion strains (JWD3, JCM290) were grown in the presence of 0.05% [w/v] arabinose and BamA/BamD were depleted by growth in 0.05% fructose. Plasmid mediated expression of proteins were induced using a final concentration of 0.5 mM IPTG or 0.2 µg/ml of anhydrotetracycline for T7 and pASK vectors, respectively.

**Table 1.**
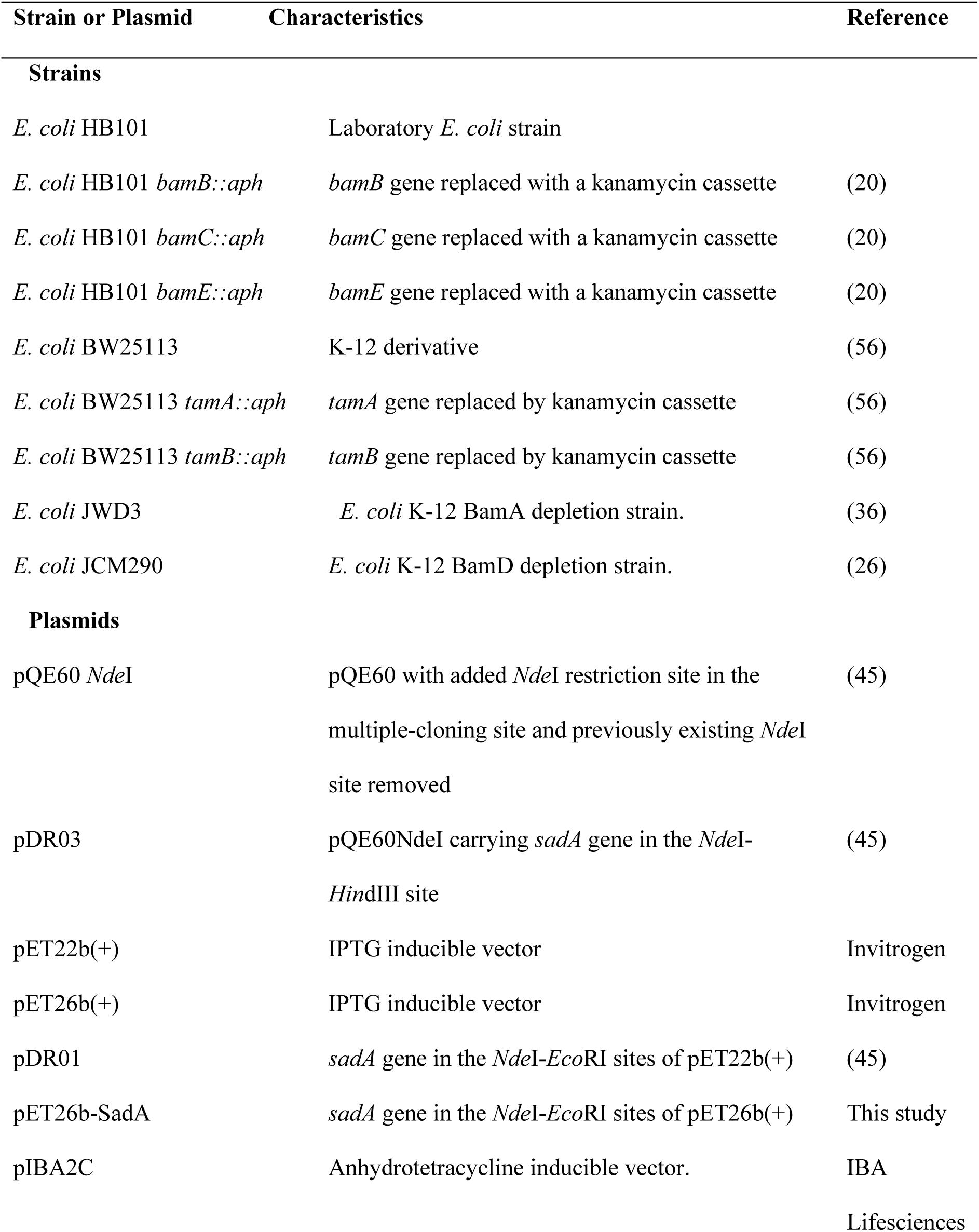
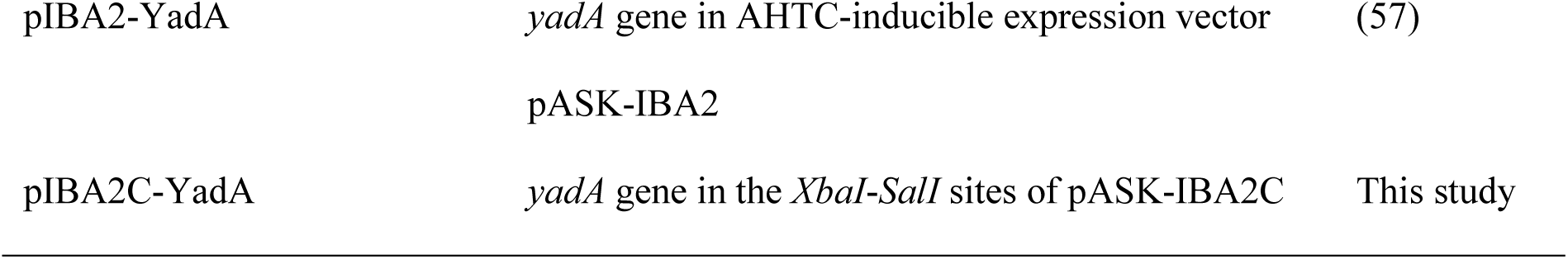
Strains and plasmids used in this study.

### DNA manipulations and genetic techniques

All plasmids were transformed into the wild-type and mutant strains via heat shock. To construct the pET26b-SadA plasmid, the *sadA* gene was *EcoR*I, *Nde*I digested from pET22b-SadA and religated into a *EcoR*I, *Nde*I digested pET26b(+) vector and the correct clone (pET26b-SadA) confirmed by sequencing. The *yadA* gene was excised from pIBA2-YadA using *XbaI* and *SalI* and re-ligated into pASK-IBA2C (IBA Lifesciences) using the same restriction enzymes to create pIBA2C-YadA.

### Protein manipulation techniques

Whole cell protein fractions were prepared by centrifuging 1 ml of overnight bacterial culture and resuspending the cell pellet in 2 x Laemmli sample buffer (Sigma) and boiled for 5 min at 100°C. Outer membrane protein fractions were prepared as previously described (38). Briefly, 50 ml of cultures were grown to an OD_600_ of 1. The pellet was resuspended in 20 ml 10 mM Tris-HCl pH 7.4 buffer and sonicated to lyse cells followed by centrifugation. The supernatant was harvested and centrifuged at 50,000 x *g*. The resulting pellets were washed in 10 mM Tris-HCL pH 7.4, containing 0.2% [v/v] Triton X-100 and subsequently centrifuged at 50,000 x *g*. The pellets, representing outer membrane fractions, were washed three times in 10 mM Tris-HCl pH 7.4 before final resuspension in 100 µl 10 mM Tris-HCl pH 7.4. Cellular fractions were analysed by SDS-PAGE and stained with Coomassie blue or transferred to a nitrocellulose membrane for Western blotting as described previously (39). Western blotting was performed with either a 1:1,000 dilution of anti-SadA antibody, 1:5,000 anti-YadA, anti-BamA, B, C, D or E antibody or 1:10,000 anti-OmpF. Finally, a 1:10,000 anti-rabbit dilution of IgG alkaline phosphatase was used before detection with NBP-BCIP (nitroblue tetrazolium chloride–5-bromo-4-chloro-3′-indolylphosphate; Sigma-Aldrich) as the substrate.

### Aggregation assay

Aggregation of the strains was determined using an aggregation settling assay as described previously (40). Briefly, overnight cultures were diluted 1:100 and grown in LB until OD_600_ of 0.8, at which point anhydrous tetracycline was added. The cultures were grown and standardised to a final OD_600_ of 1.0 and mixed vigorously for 15 s prior to the start of the assay. Samples (150 µl) were removed at 30 min intervals taken approximately 0.1 cm from the liquid surface and transferred into a microtitre plate maintained on ice. At the end of the experiment the OD_600_ values were measured using a microtitre plate reader.

### Biofilm assay

The ability of strains to form biofilms was determined using the microtitre plate biofilm assay as described previously (41). Briefly, overnight cultures were inoculated into fresh M9 minimal media to an OD_600nm_ of 0.02. The plate was incubated overnight at 37°C with shaking. Following incubation, the cultures were removed from the plate and 150 µl of 0.1% [w/v] crystal violet stain was added to each well. Upon removal of the stain, the plates were washed with distilled water and left to dry. Addition of an 80:20 ethanol: acetone solution solubilised the crystal violet, and the OD_620nm_ was measured. Each experiment was performed with 12 technical replicates and three biological replicates. Significance was determined by students *t*-test.

### Bacterial adherence to extracellular matrix (ECM) molecules

Bacterial adherence to ECM molecules was measured via growth essentially as described previously (39). Briefly, cells from overnight cultures were grown to OD_600_ of 1 and appropriately induced for two hours. These cells were washed in PBS, resuspended to an OD_600_ of 1 and added to wells coated with 20 μg/ml of collagen I (Sigma-Aldrich). BSA-coated wells (50 μg/ml) were used as a control. After 1 h incubation the wells were washed to remove non-adherent bacteria. Adherent cells were incubated in LB for a further 5 h prior to enumeration by direct colony counting on agar plates.

### Immunofluorescence and microscopy imaging

Fixation and preparation of bacterial cells for live cell imaging was performed essentially as previously (42). Briefly, fixed cells were put onto poly-L-lysine-coated coverslips, washed three times with PBS and then blocked for 1 h in PBS containing 1% [v/v] bovine serum albumin (Europa Bioproducts). Coverslips were incubated with 1:500 anti-YadA antibody for 1 h, washed three times with PBS, and incubated for an additional 1 h with Alexa Fluor® 488 goat anti-rabbit IgG. The coverslips were washed a further three times with sterile PBS before mounting onto glass slides and visualized using either phase contrast or fluorescence using Leica DMRE fluorescence microscope (100× objective) -DC200 digital camera system.

## RESULTS

### BamA and BamD are required for the secretion of SadA

Previously, we determined that BamA was required for the biogenesis of the trimeric autotransporter YadA (36). To determine if this observation was true for other TAAs, we investigated whether BamA was essential for the biogenesis of a sequence divergent TAA, namely SadA. An expression vector encoding full-length SadA (pDR03) was transformed into the BamA depletion strain *E. coli* JWD3 (36). After growth in the presence or absence of arabinose, outer membrane fractions were harvested and SadA was detected by Western immunoblotting. SadA could be detected in wild-type strains of *E. coli* and in *E. coli* JWD3 under BamA replete conditions (Fig. 1A). In contrast, like BamA and the BamA-dependent protein OmpF, SadA levels were severely diminished after growth in BamA-depleted cells.

**Figure 1.**
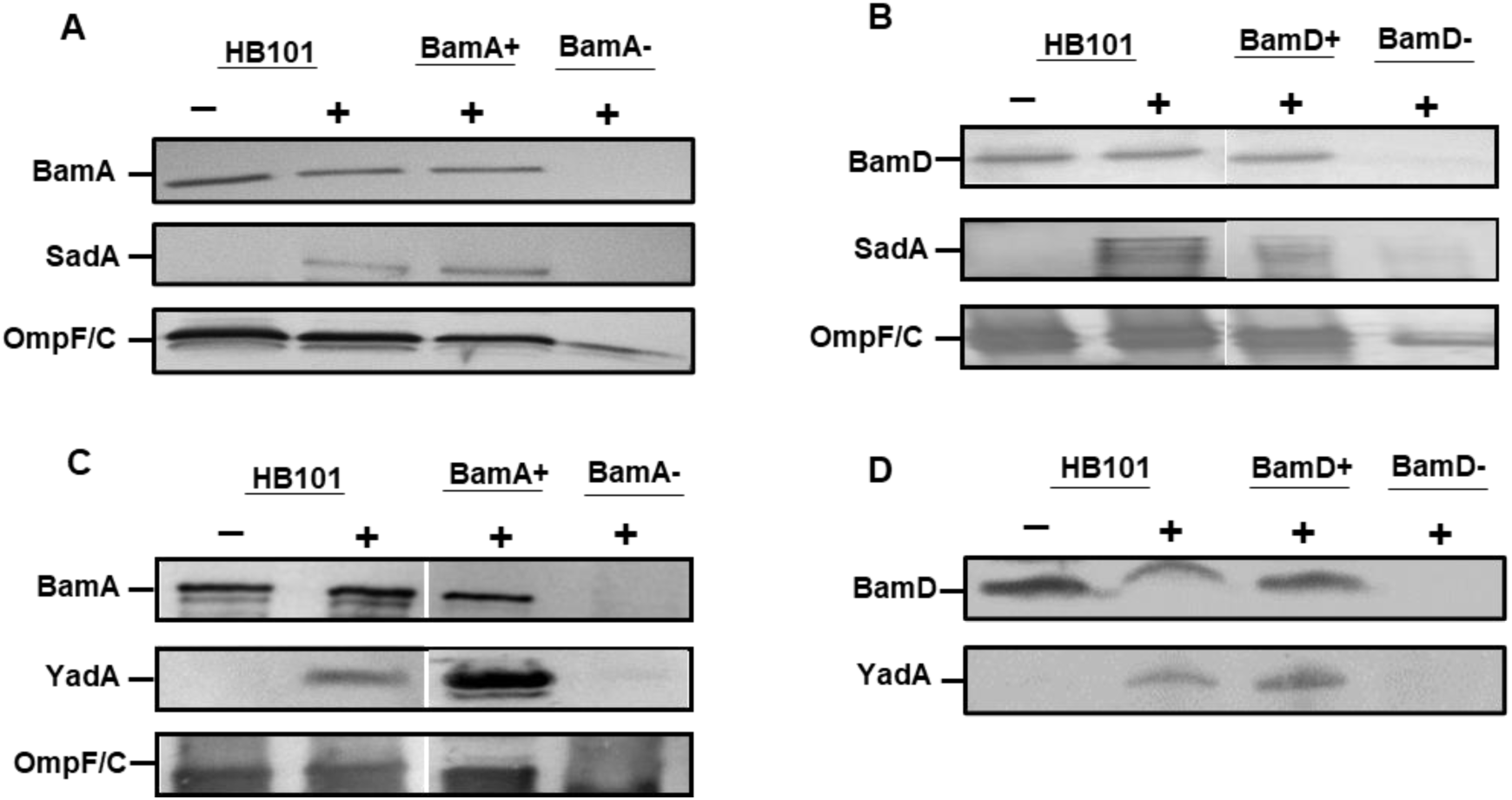
BamA and BamD are required for secretion of SadA. (A) Western blot analysis of outer membrane fractions of HB101 and BamA depletion JWD3 strain (BamA+/- depletion) with either empty vector (-) or pET22b-SadA (+) plasmids. Fractions were probed with anti-BamA, anti-SadA, and anti-OmpF/C antisera. Bands indicate the presence of the protein in the outer membrane. (B) Similar analysis of HB101 and BamD depletion JCM290 strain (BamD+/- depletion) with EV or SadA+ plasmids. Fractions are probed with anti-BamD, anti-SadA, and anti-OmpF/C antisera. (C) Western blot analysis of outer membrane fractions of HB101 and JWD3 depletion strain (+/- BamA) with either empty vector (-) or YadA (+). (D) Western blot analysis of JCM290 (+/- BamD) with either empty vector (-) or YadA (+).

BamD is the second component of the Bam complex that is vital for cell viability (26). To determine whether BamD is required for TAA secretion we followed a similar method to BamA above. The secretion of SadA to the outer membrane was investigated in a BamD depletion strain (JCM290). When BamD is depleted from the cell, the protein levels of SadA and OmpF/C are considerably depleted in the outer membrane (Fig. 1B), which was similar to the results observed for BamA.

To further confirm the observations that the essential Bam components are required for the secretion of TAAs, YadA was investigated for secretion under BamA and D depletion conditions. HB101, JWD3 and JCM290 were transformed with an expression vector that expressed *yadA* upon addition of anhydrotetracycline. After growth in the absence or presence of arabinose, the outer membrane proteins were isolated as previously described. The proteins were analysed by SDS-PAGE and Western immunoblot. Under BamA replete conditions, YadA was detected in the outer membranes of both HB101 and JWD3. Upon depletion of BamA, YadA was no longer detectable in the outer membrane (Fig. 1C), as previously reported (36). Similarly, when BamD was depleted from the cells, YadA could no longer be detected in the outer membrane (Fig. 1D). These data reinforce previous observations that BamA is required for the biogenesis of TAAs and demonstrate for the first time that BamD is essential for TAA biogenesis.

### The non-essential Bam components are not required for outer membrane localisation of TAA’s

While BamA and BamD are required for the biogenesis of TAAs, we sought to investigate the contribution of the nonessential components of the Bam complex (BamB, C, and E) for TAA biogenesis. *E. coli* wild-type and *bamB, bamC*, and *bamE* deletion mutants (20) were transformed with pDR03 (SadA+) or pIBA2-YadA (YadA+) or empty vector. Outer membrane protein fractions of each sample were probed for the presence of SadA, YadA and each Bam component. The Bam mutants were confirmed by Western blot analysis (Fig. 2A) and strains expressing either SadA or YadA were shown to have the protein present in the outer membrane fractions (Fig. 2B). The absence of BamB, BamC, or BamE did not affect the accumulation of either YadA or SadA in the outer membrane (Fig. 2B).

**Figure 2.**
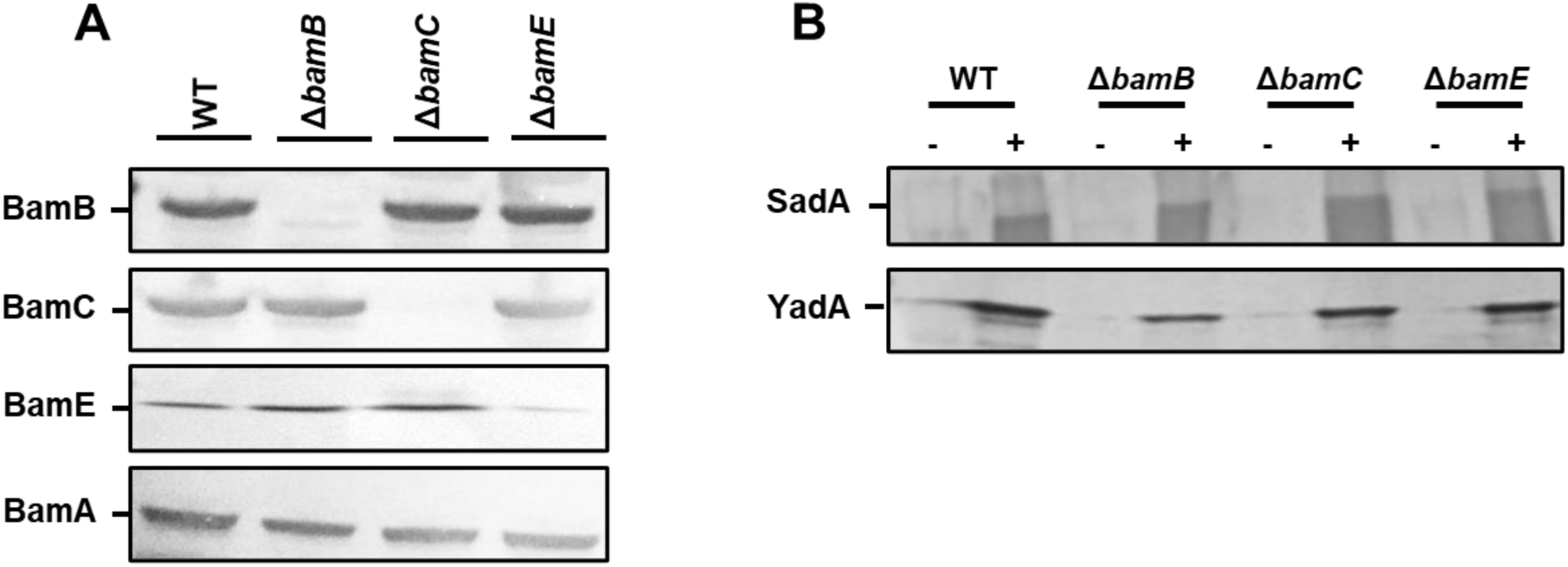
Non-essential BAM components are not required for outer membrane localisation of trimeric autotransporters. (A) Western blot analysis of wild-type HB101 and *bamB, C* or *E* mutant strains confirming the loss of each protein in the corresponding mutant. Outer membrane fractions were probed with either anti-BamA, B, C, E specific antibodies. (B) Outer membrane fractions of wild-type and bam mutant strains expressing either an empty vector control (-) or SadA/Yada (+) analysis by Western blot. Protein fractions were probed using either anti-YadA or anti-SadA specific antibodies confirming the presence of the proteins in the outer membrane.

### The Tam complex is not essential for outer membrane localisation of TAA’s

Earlier studies demonstrated that the Tam complex is required for the correct biogenesis of members of the T5aSS. Therefore, we hypothesised that the TAAs proteins would also require the assistance of the Tam complex for biogenesis. To test this, we first created *tamA* and *tamB* null mutants and confirmed by Western blot analysis of whole cell protein fractions (Fig. 3A). The resulting mutants were transformed with either the empty vector or plasmids encoding either SadA or YadA and outer membrane fractions probed for presence of YadA or SadA (Fig. 3B). In the absence of TamA or TamB, both YadA and SadA are incorporated into the outer membrane at levels similar to the wild-type parental strain, suggesting that the Tam complex is not essential for the localisation of TAAs to the OM (Fig. 3B).

**Figure 3.**
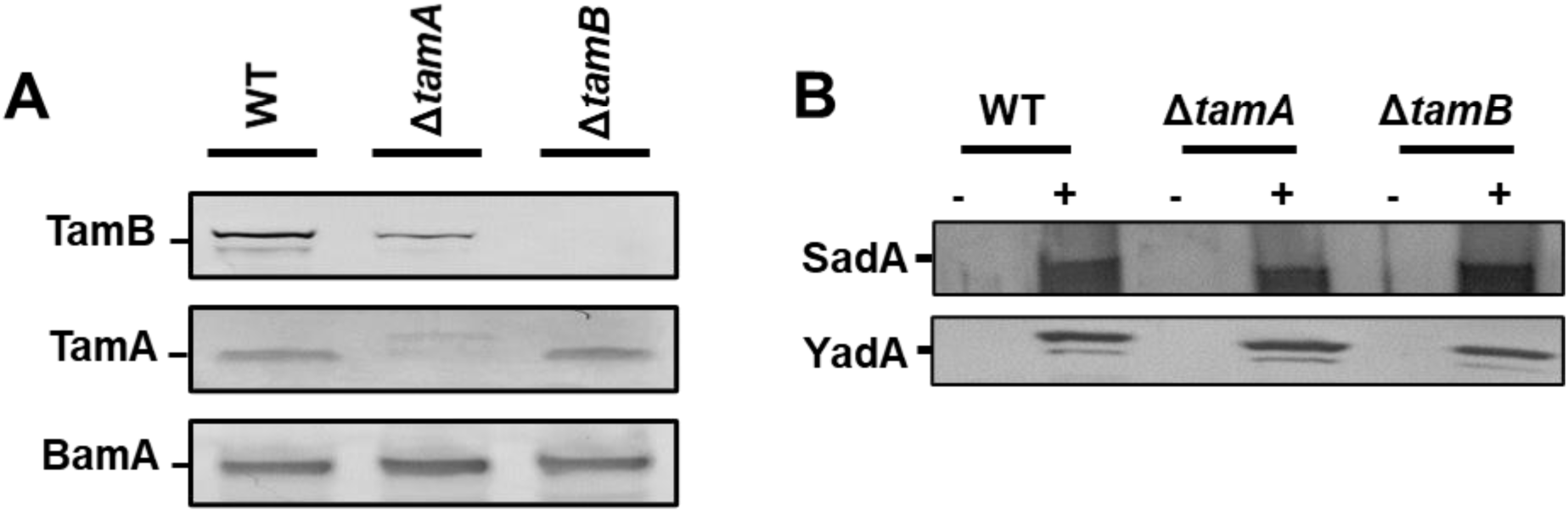
TamA and TamB are not required for outer membrane insertion of trimeric autotransporters. (A) Western blot analysis of wild-type HB101 and *tamA* and *tamB* mutant outer membrane fractions confirming the loss of each protein in the corresponding mutant using antibodies specific to TamA and B with anti-BamA specific antibodies used as a positive control. (B) Western blot analysis of wild-type HB101 and *tamA* and *tamB* mutant outer membrane fractions expressing either empty vector (-) or YadA/SadA (+) using YadA or SadA specific antibodies to confirm the presence of the proteins in the outer membrane.

### TAA passenger domains are secreted to the cell surface in *bam* and *tam* mutants

Previously, we observed intermediates of the T5aSS that are localised to the outer membrane but are unable to translocate their cognate passenger domains to the cell surface (43). Therefore, we hypothesised that whilst the non-essential Bam components and the Tam components were not required for localisation to the outer membrane, they might be responsible for holding the C-terminal translocation unit and/or passenger domains in a conformation that permits secretion of the passenger domain to the cell surface. The surface exposure of YadA was determined by specific immunofluorescence microscopy in WT, BamB, C, E and TamA and B mutants expressing YadA (Fig. 4). YadA was detectable on the surface of all strains containing the YadA vector including all three Bam mutants and both Tam mutants (Fig. 4).

**Figure 4.**
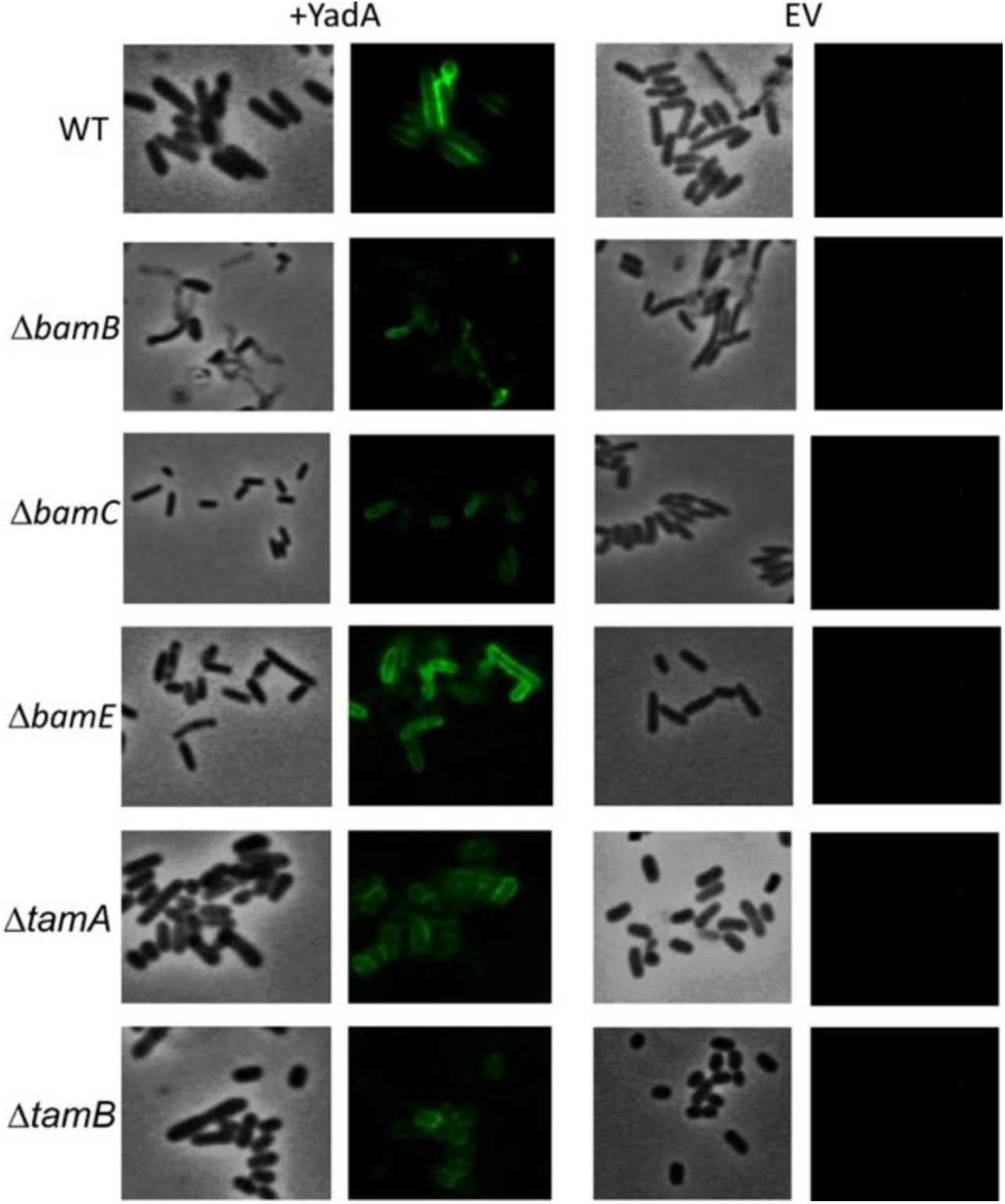
Bam and Tam components not required for OM localisation of trimeric autotransporters. Immunofluorescence imaging of wild-type, and *bamB, C, E, tamA* and *B* mutants PFA fixed cells containing either empty vector (EV) or YadA plasmid (+YadA) probed with anti-YadA antisera. The figure shows light microscope and fluorescent images of cells carrying empty vector (EV) or expressing *yadA* (+Yada).

### TAA passenger domains are functional in *bam* and *tam* mutants

Although the Tam and the non-essential Bam components are not required for surface localisation of the passenger domain, they may be required for the correct folding and function of TAA passenger domains on the cell surface. Thus, we investigated whether the TAA proteins retained functional competence in these mutant backgrounds. As YadA is known to mediate binding to collagen I and to promote autoaggregation (44), the *E. coli bamB, bamC* and *bamE* mutants expressing *yadA* were tested in a growth-based collagen binding assay. Expression of *yadA* in WT and *bamC* and *E* mutant strains promoted a significant increase in binding to collagen I compared to the empty vector controls (Fig. 5A). Whilst there was an increase in bacterial adherence to collagen I in the *bamB* mutant, this was not significantly different to the empty vector control (Fig. 5A). As *bamB* mutants are known to have a reduced growth rate, we concluded that this lack of significance was probably due to slower growth of the *bamB* mutant. YadA is also known to promote cell-cell autoaggregation therefore we tested the ability of YadA to promote autoaggregation in the non-essential Bam mutants. Induction of *yadA* expression in all strains led to significant autoaggregation, in contrast to empty vector control strains which had no discernible aggregation (Fig. 5B). Previously, we demonstrated that SadA could promote biofilm production *in vitro* (45). Thus, we investigated the ability of wild-type and Bam mutant strains, either expressing or not expressing SadA, for their ability to form a biofilm on polystyrene plates when grown in M9 minimal medium. All strains expressing SadA had significantly (*P* < 0.05) higher biofilm formation than the empty vector controls (Fig. 5C).

**Figure 5.**
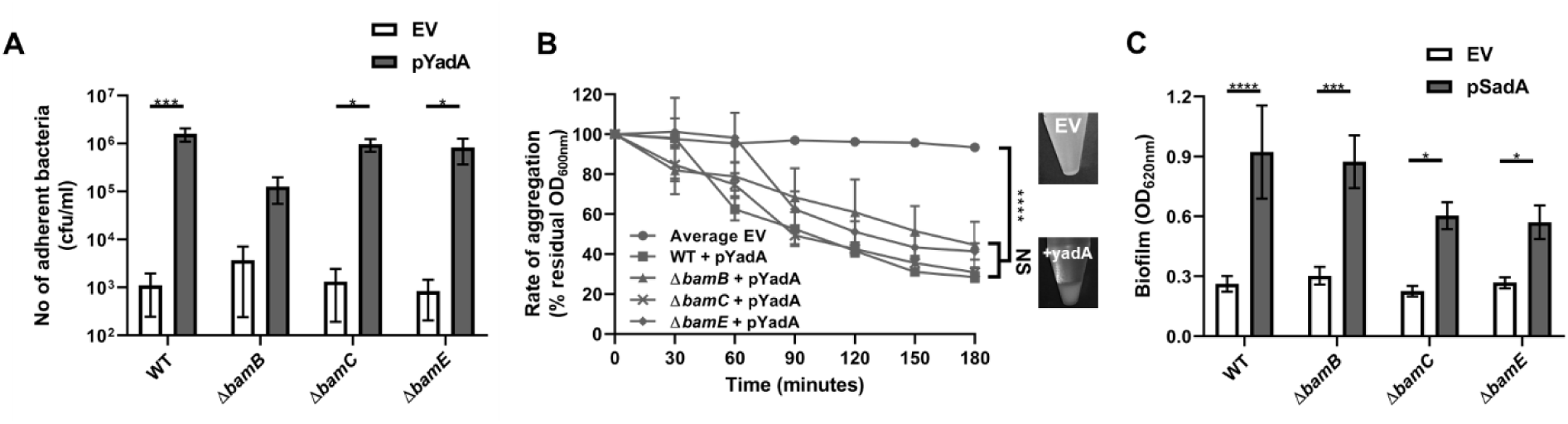
Trimeric autotransporters are functional in the absence of the non-essential Bam components. (A) Collagen I binding by wild-type and Bam mutant HB101 strains containing either EV or YadA+ plasmid. Adhesion was examined via measuring growth in microtitre plates after binding and washing. Significance (p<0.05) was determined by a Students *t*-test. Error bars represent standard error of the mean (SEM) of at least 8 technical replicates and 3 separate experiments. (B) Aggregation of wild-type and mutant HB101 strains containing either EV (empty vector) or YadA+ plasmid. The pellet formed by autoaggregation can be visibly compared after three hours (insets). Significance (p<0.05) was determined by a 2-way Anova with Turkey’s correction for multiple comparisons. Error bars represent SEM of 3 separate experiments. (C) Biofilm formation by wild-type and mutant HB101 strains containing either EV or SadA expressing plasmid. Biofilm formation was examined in polystyrene microtitre plates. Significance (p<0.05) was determined by a Students *t*-test Error bars represent standard deviation of at least 12 technical replicates and 5 separate experiments. Significance was represented as follows: * p<0.05, ** p<0.01, *** p<0.001, and **** p<0.0001.

To investigate whether TamA and TamB were required for correct folding of TAA passenger domains, the *tam* mutants described above were monitored for SadA and YadA function. As above we tested the WT and *tamA* and *B* mutants for their ability to bind collagen I. All three strains expressing YadA had significantly higher collagen I binding compared to empty vector controls (Fig. 6). We found that *tamA* and *B* mutants in *E. coli* form high biofilm (data not shown) and thus were unable to determine whether SadA could promote biofilm formation in these strains. These data suggest that TAAs are secreted, folded and function even in the absence of BamB, C, E, TamA or B mutants.

**Figure 6.**
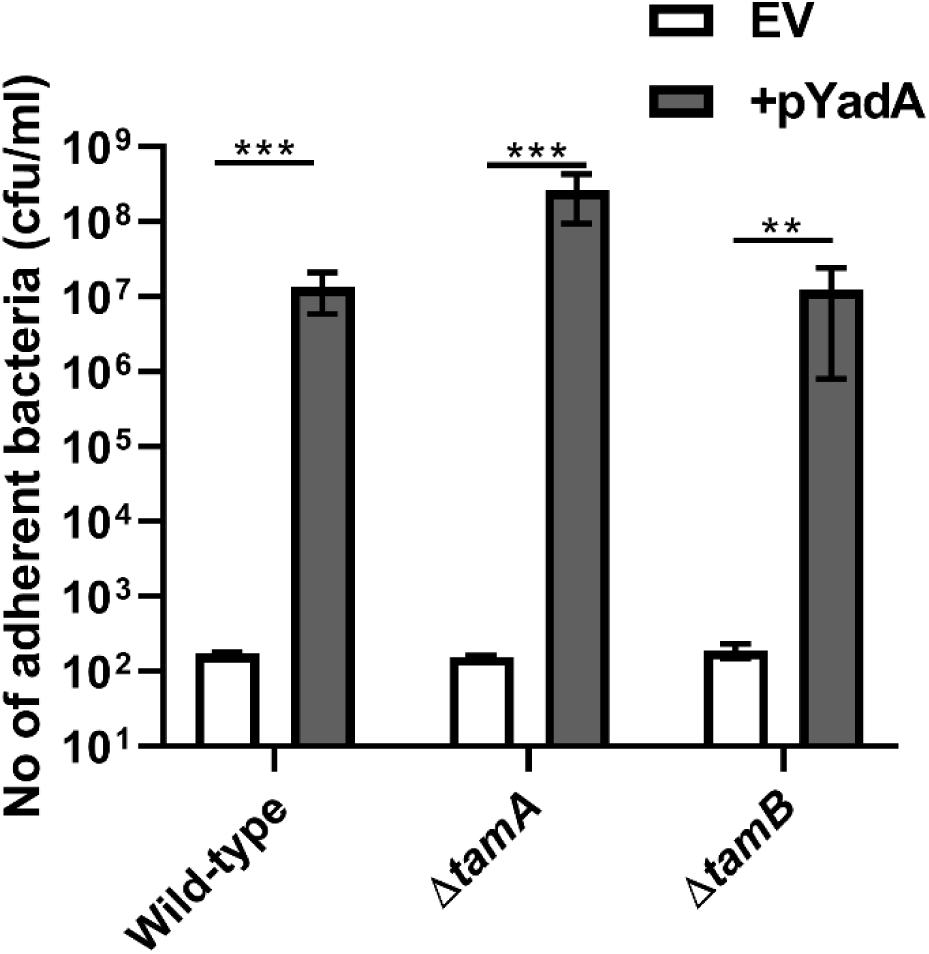
YadA secreted in the absence of TamA and B is able to bind collagen I. (A) Collagen I binding by wild-type (WT) and mutant *tamA* and *tamB* strains containing either empty vector (EV) or the YadA+ plasmid. Adhesion was examined by measuring growth in microtitre plates after binding and washing. Significance (p<0.05) was determined by a Students *t*-test. Error bars represent standard deviation of 3 biological replicates. Significance was represented as follows: ** p<0.01 and *** p<0.001.

## Discussion

The proteins SadA and YadA are both trimeric autotransporter adhesins, however they vary greatly in sequence homology, length, and function. YadA is the prototypical TAA, its trimeric structure composed of a β-barrel in the outer membrane, with a single stalk, neck and head domain extending 35 nm into the extracellular environment (46). In contrast, SadA extends 108 nm and is composed of multiple stalk, neck and head domains alternatingly dispersed along the length of the passenger domain (47). Here we found that both BamA and BamD are required for the secretion and function of YadA and SadA, and by implication all TAAs. This is in agreement with studies on other outer membrane proteins (20), and is further evidence that BamA and BamD work in unison to assemble proteins into the outer membrane (26, 48). In contrast, both SadA and YadA were functionally assembled in the outer membrane in *bamB, bamC* and *bamE* mutant strains. These three lipoproteins have been shown to be non-essential in *E. coli* and although they are needed for the secretion of some outer membrane proteins (48, 49), are also not essential for others (50-52). In addition, and consistent with previous studies (37), we observe that TamA and B are not required for the biogenesis or function of TAA proteins. However, TamA and B have been implicated in the secretion of classical autotransporters (27), suggesting TamA and B may be involved in secretion of a specific subset of Type 5 proteins or may have importance under specific conditions.

Recent evidence suggests that trimerisation of TAA’s occurs in the periplasm prior to outer membrane insertion (54) and that TAA passenger translocation occurs via hairpin intermediates (55). The ability of SadA to induce biofilm formation and YadA to induce autoaggregation in all of the mutants at levels similar to the wild-type suggests that these proteins are functionally assembled in a folded state on the surface of the bacterium. These data indicate that the non-essential Bam components and the Tam components are not essential for the translocation or folding of TAAs on the bacterial cell surface. This is in agreement with other studies that demonstrate other proteins such as Pet, TolC are functionally assembled in *bamB* mutants (20, 49). Additionally, TAA proteins such as NhhA exist in bacterial species (*Neisseria meningitidis*) that do not have a BamB homolog, suggesting it is not required for proper assembly of this protein class (51). Interestingly, although SadA protein levels in outer membrane preparations were similar in all strains, biofilm formation induced by SadA was significantly lower in a *bamC* and *bamE* knockout compared to wild-type (p<0.05) (Fig. 5C). This suggests that these lipoproteins may play a more subtle role in TAA secretion, and multiple lipoprotein knockouts may lead to secretion defects. Indeed, in *N. meningitidis* a *bamC/E* double knockout leads to a lethal phenotype (51) and it has recently been shown that a double *bamB*/*bamE* mutant is lethal in *E. coli* (53), suggesting the non-essential lipoproteins have some redundancy.

In conclusion, we show that the non-essential components of the Bam complex and both TamA and B are not essential for TAA assembly. Given the complex nature of TAA biogenesis, due to large and complex passenger structures and the need for trimerisation, it is plausible that loss of Bam and/or Tam proteins could lead to a reduced efficiency of TAA passenger translocation. Further experimental evidence would be required to investigate this hypothesis.

